# Astrocyte plasticity ensures continued endfoot coverage of cerebral blood vessels and integrity of the blood brain barrier, with plasticity declining with normal aging

**DOI:** 10.1101/2021.05.08.443259

**Authors:** William A. Mills, Shan Jiang, Joelle Martin, AnnaLin M. Woo, Matthew Bergstresser, Ian F. Kimbrough, Harald Sontheimer

## Abstract

Astrocytes extend endfeet that enwrap the vasculature. Disruptions to this association in disease coincide with breaches in blood-brain barrier (BBB) integrity, so we asked if the focal ablation of an astrocyte is sufficient to disrupt the BBB. 2Phatal ablation of astrocytes induced a plasticity response whereby surrounding astrocytes extended processes to cover vascular vacancies. This occurred prior to endfoot retraction in young mice yet occurred with significant delay in aged animals. Laser-stimulating replacement astrocytes showed them to induce constrictions in pre-capillary arterioles indicating that replacement astrocytes are functional. Inhibition of EGFR and pSTAT3 significantly reduced astrocyte replacement post-ablation yet without perturbations to BBB integrity. Identical endfoot replacement following astrocyte cell death due to reperfusion post-stroke supports the conclusion that astrocyte plasticity ensures continual vascular coverage so as to retain the BBB. Together, these studies uncover the ability of astrocytes to maintain cerebrovascular coverage via substitution from nearby cells and may represent a novel therapeutic target for vessel recovery post-stroke.

## Introduction

Astrocytes serve essential roles in supporting normal brain physiology (1). This is made possible, in part, by the extension of large, flattened processes, called endfeet, that wrap around blood vessels. Thought to cover up to ~99% of the cerebrovascular surface (2), astrocytic endfeet, in conjunction with pericytes (3), help to maintain expression of molecules that form the blood-brain barrier (BBB)-including endothelial tight junction, enzymatic, and transporter proteins (4–6). Astrocytic endfeet also mediate neurovascular coupling, also known as functional hyperemia, whereby local blood flow adjusts to local energy demand. Astrocytes sense changes in neuronal activity via purinergic receptors that cause increases in [Ca^2+^]_i_, leading to the release of vasoactive molecules onto pericytes at capillaries (7–8), or directly release vasoactive signals onto arterioles (9–12), leading to change in vessel diameter.

Interestingly, a number of CNS diseases are marked by retraction or separation of astrocytic endfeet from blood vessels-a phenotype often simultaneously presenting with vascular deficits such as altered blood-brain barrier permeability or elevated CSF-to-serum albumin ratio, which is indicative of blood-brain barrier breakdown. Examples include multiple sclerosis (13), major depressive disorder (14–16) ischemia (17–19), and even normal biological aging (20–22). We previously demonstrated separation of endfeet from the vasculature due to invading glioma cells (23) as well as due to amyloid accumulation on vessels (24). Both conditions resulted in disruption to neurovascular coupling and, in the case of glioma, BBB breakdown. This raises the question of whether astrocyte endfeet are required to maintain an intact BBB, or whether lost endfeet can be replaced by other astrocytes as has been shown for pericytes (25). Moreover, since changes in astrocyte morphology and function are known to occur with physiological changes of the organism-i.e. parturition, lactation, chronic dehydration, starvation, voluntary exercise or sleep deprivation (26–29) – it is possible that astrocyte association with blood vessels is equally dependent on the physiological context.

Given the multitude of conditions marked by regions of abnormal vasculature lacking endfoot coverage, we were interested in determining whether replacement endfeet have functional relevance in maintaining blood-brain barrier integrity and astrocyte-vascular coupling. Using multiphoton imaging through a cranial window, we were able to induce single-cell apoptosis using the 2Phatal method (30) to question whether loss of endfeet on blood vessels would be compensated for by neighboring cell(s). We find remarkable plasticity, discovering that the ablation of single astrocytes reliably causes innervation by neighboring cells. In young animals, this happens before the ablated cell completely retrieves its process; yet in 12-month-old animals, replacement occurs with a significant 1-2h delay after the ablated cell has vacated the vessel. Endfoot replacement engages the EGFR/STAT3 signaling pathway as pharmacological inhibition via AG490 injection impairs replacement. Once in place, the replacement endfeet have the ability to vasoconstrict precapillary arterioles like normal astrocytes. Despite recent evidence that global astrocyte loss results in impairments of the bloodbrain barrier (31), we did not find this to be the case even when replacement endfoot coverage was impaired by inhibition of EGFR and/or STAT3 phosphorylation. Finally, we demonstrate using focal photothrombosis that astrocyte apoptosis following reperfusion triggers a focal gliovascular plasticity response wherein astrocyte-vascular coverage is maintained. Together, these results reveal a novel process in which astrocytes cover for neighboring cells to maintain vascular coverage in an EGFR/pSTAT3-dependent manner.

## Materials & Methods

### *In vivo* multiphoton imaging through a cranial window

All surgeries were performed as described previously (19–20) with slight modifications. Following induction of surgical plane anesthesia with 2-5% isofluorane, pre-operative analgesics and antibiotics were administered intraperitoneally. Following this, the hair and skin of the skull was removed, and a 3×3 mm craniectomy anterior to lambda and posterior to bregma was subsequently performed on one hemisphere. A durotomy was performed next, followed by placement of a 3×3mm #1 cover glass that was then affixed and sealed with dental cement. All mice were allowed to recover for 5-7 days before experiments commenced. For imaging, animals were placed on a Kopf stereotax with heating pad. While imaging, animals were lightly anaesthetized (~100 beats per minute), and their vitals constantly monitored. Cerebral vessels were visualized by retro-orbital injection of 70kDa TRITC, 3kDa TRITC, and/or 967 Da Cadaverine Alexa Fluor 555. A Chameleon Vision II(Coherent) laser tuned to 870nm was used to excite all dyes. Optical sections were acquired using a four-channel Olympus FV1000MPE multiphoton laser scanning fluorescence microscope equipped with a XLPLN25X/1.05 NA water-immersion objective (Olympus). Z projections were created using FIJI(NIH) and NIS-Elements(Nikon).

### 2Phatal Ablation

To induce single-cell apoptosis in astrocytes, Hoechst 33342 (ThermoFisher catalogue number H5370) was applied topically (0.04 mg ml^-1^ diluted in PBS) to the durotomized cortex of Aldh1l1-eGFP mice for 10 minutes and washed thoroughly with cold 1XPBS.-To ablate, an 8×8 μm square ROI was placed over dual eGFP Hoechst positive astrocyte nuclei whose soma was either on the vessel or that had endfeet contacting the vasculature. Pixel dwell time was set to 100 μs/pixel, laser wavelength was set to 775 nm and photobleaching was achieved by scanning for a duration of 20 s. A Newport Model 1919-R power meter with a silicone based OD3 photodetector attached was used to determine power at the objective for all ablation experiments, and a range of 2.33 mW to 53.4 mW was used for all experiments, where power increased with increasing depth and/or decreasing Hoechst intensity measured in the activation ROI. eGFP was visualized by setting laser wavelength to 870 nm.

### Quantification of astrocyte morphometrics

To determine the number of astrocytes extending processes to innervate vascular vacancies, NIS elements was used to compare the baseline z-stack to z-stacks from time points following complete removal of ablated astrocytes. Specifically, ablated astrocytes and the optical section(s) their soma occupied was denoted in the baseline image. The surrounding vascular landmarks and astrocytes that weren’t ablated could therefore serve as fiduciary landmarks when evaluating that same field post-ablation. Any astrocyte at post-ablation timepoints that appeared to extend processes and innervate a vascular vacancy was identified in the baseline image, again using 1) the surrounding vascular profile, 2) astrocytes that were not ablated, and 3) z location of the astrocyte in question relative to the ablated astrocyte. If, at baseline, the replacement astrocyte in question did not appear to have processes interacting with the vasculature, even upon dramatically increasing the look up table, it was considered a replacement astrocyte. The total numbers of cells fulfilling these criteria were reported as the total number of replacement cells.

For analysis of replacement kinetics, only astrocytes extending clear processes to the vascular interface were chosen for ablation. Longitudinal imaging was performed following astrocyte ablation. Specifically, images were acquired twice per day to determine if the abated astrocyte’s soma has begun to swell, indicative that it would undergo phagocytosis within the next 24 hours. Once identified, z stacks were captured every 5 minutes until the time point of first seeing a process from a replacement astrocyte occupy the vacant vascular territory or identifying the time of fluorescence fading in the process of the ablated astrocyte. To quantify the number of minutes to endfoot replacement in 4-month and 12-month-old mice, the time of fluorescence fading in the process of the ablated astrocyte was considered time point zero. From there, the number of minutes until observing a process from a replacement astrocyte occupy the vascular territory of the ablated was then determined from the captured z-stacks. If the replacement process made contact with the vessel prior to time point zero, this was reported as negative minutes. If the replacement process made contact with the vessel after time point zero, this was reported as positive minutes.

For eGFP volumetric analysis in AG490 studies, images were opened in NIS elements volume viewer and underwent background subtraction using the rolling ball radius feature. A median filter and binary threshold were subsequently applied and eGFP volume recorded. The same number of optical sections were used for time points being directly compared. To ensure that the region of astrocyte ablation was compared pre- and postablation, the rotating rectangle feature was used. This allows for area selection in an image without changing the underlying metadata. An ROI of the same size was applied to images of time points being compared. so that it could be used as size reference for the rotating rectangle.

#### Quantification of BBB leakage

Prior to astrocyte ablation (time point 1), a 30-μL bolus of 100mg/ml 3kDa TRITC (D3307 Invitrogen) was retro-orbitally injected, and a timer started at the moment of injection. The mouse was moved immediately to the stereotax under the microscope, and a z stack was captured. The times at the start of image capture and end of image capture were recorded to enable proper comparison at all subsequent time points. This process was repeated at the moment of fluorescence fading in the process at the vascular interface (time point 2) and the same imaging parameters used as that of the baseline image. FIJI (ImageJ) was used to create sum intensity projections and the same number of optical sections was used for all time points, where optical sections in which the ablated astrocyte was covering the vasculature were selected. Background subtraction was performed using the rolling ball radius feature, and an average fluorescent value was measured at the location of endfoot coverage at both time points. For the induction of vessel injury as a positive control, 870nm line scans at a laser power of 50 to 85mW were applied across the vessel wall for 90 to 120 seconds. As in prior BBB measurements, a z stack was captured at the same time after retro-orbital injection of 3kDa TRITC, and the average intensity just beside the damaged vessel was compared at both time points, pre- to post-vessel injury.

#### *In vivo* replacement astrocyte induced-precapillary arteriole constriction

To determine if replacement astrocytes can vasoregulate precapillary arterioles, Aldh1l1cre x GCaMP5G mice received a cranial window following the methodology described above. The first branching capillary segment from Alexa 633 hydrazide positive penetrating arterioles was selected for imaging if the soma of astrocytes making contact with the precapillary arterioles were on a focal plane similar to the vessel. Using a four-channel Olympus FV1000MPE multiphoton laser scanning fluorescence microscope equipped with a XLPLN25X/1.05 NA water-immersion objective (Olympus), single-plane images of 1024×800 pixels were obtained every 3 seconds. Astrocytes were targeted for laser irradiation by selecting a focal plane where the soma and associated precapillary arteriole were visible. Astrocyte stimulation was achieved using a 4μm^2^ circular region of interest centered within the astrocyte soma for 800 milliseconds at 7-10x imaging power levels. A two-minute measurement was recorded before and after astrocyte activation. Vessel diameter was measured as the cross-section of the vessel using FIJI (ImageJ) software. Motion correction in videos was performed using the Intravital Microscopy Toolbox ImageJ macro developed by Soulet et al (32).

#### Rose Bengal Photothrombosis

To focally and transiently induce Rose Bengal intravascular clot formation, Aldh1l1-eGFP mice were retro-orbitally injected with Alexa 633 hydrazide and penetrating arterioles identified prior to the beginning of the experiment. Rose Bengal was then retro-orbitally injected, and all mice <20g received a 25μL injection whereas mice >20g received a 50μL injection. Mice were then transferred to the multiphoton as soon as possible and a square ROI was placed over a penetrating arteriole. The dimensions of the ROI were based upon the dimensions of the penetrating vessel, and imaging was conducted 90μms from the surface of the brain. Imaging duration was set to two minutes and wavelength set to 870nm. Laser power was set between 50-90 mW, with power determined on the average intensity of Rose Bengal at the beginning of the experiment. If only part of the vessel was occluded at the end of two minutes, the vessel would be imaged in laser scanning mode for until dye nucleation was complete. In all instances, this was not longer than two minutes. Upon successful intravascular clot formation, a z-stack was captured to visualize the clot, and then the animal was placed back in its home cage. The animal was then subsequently imaged one hour later to confirm that the clot had cleared. All instances of reported endfoot replacement come from depths below the plane of dye nucleation.

#### Drug Treatment

AG490 at 10mg/kg in 40%DMSO/PBS was subcutaneously injected daily for initial studies comparing percent increase in eGFP volume at day post ablation 5 relative to baseline (Figure 6). This dosage was increased to 3x/day for studies aiming to prolong the time a vessel region remained vacant post-ablation (Supplementary Figure 4).

#### Statistics

GraphPad Prism software was used to perform all statistical analyses. Details for every statistical test are reported in the figure legends. Every parametric test used was validated by first performing tests of normality on the dataset once outliers were removed. Parametric tests were further selected based on datasets having equivalent or different standard deviations, where a difference of <1.5 was counted as being equal.

## Results

### Focal ablation of single astrocytes does not breach the blood-brain barrier but induces an astrocyte endfoot replacement response

Given our previous finding of compromised blood-brain barrier (BBB) integrity in regions of focal endfoot separation due to invading glioma cells, we questioned if the focal ablation of a single astrocyte is sufficient to induce breaches in blood-brain barrier integrity. To do so, we implanted cranial windows in mice and adopted the 2Phatal ablation method developed by Hill et al (30). This technique employs the focal illumination properties of a femtosecond-pulsed laser to activate the nucleic-acid binding Hoechst dye **(Figure 1a)**, triggering apoptosis. Given the ability to induce single-cell apoptosis, we were able to image astrocytes in Aldh1l1-eGFP mice up to and beyond removal of their corpse, which included the retraction of their endfeet **(Figure 1b**). We also employed Alexa Fluor 633 hydrazide (**Supplementary figure 1a-b)** to selectively label arterioles (33) so as to avoid perturbations to pericyte physiology, which are known to play a role in BBB integrity and vasodilation of capillaries. Comparing the extravasation of retro-orbitally injected 3kDa TRITC at the time of endfoot retraction to baseline revealed no apparent difference in BBB permeability **(Figure 1c-e**). 3kDa TRITC was chosen because initial attempts to use the ~1kDa Cadaverine revealed that at baseline, this dye extravasates and is taken up by astrocytes **(Supplementary Figure 2a)**. Positive control experiments utilizing direct laser irradiation of the vasculature demonstrated that we were able to detect leakage of 3kDa TRITC from the vasculature **(Supplementary Figure 2c-g)**. At the initial stages of astrocyte endfoot retraction, we observed nearby neighboring astrocytes extended processes to the soon-to-be vacancy left by the ablated astrocyte **(Figure 1e)**.

**Figure 1.**
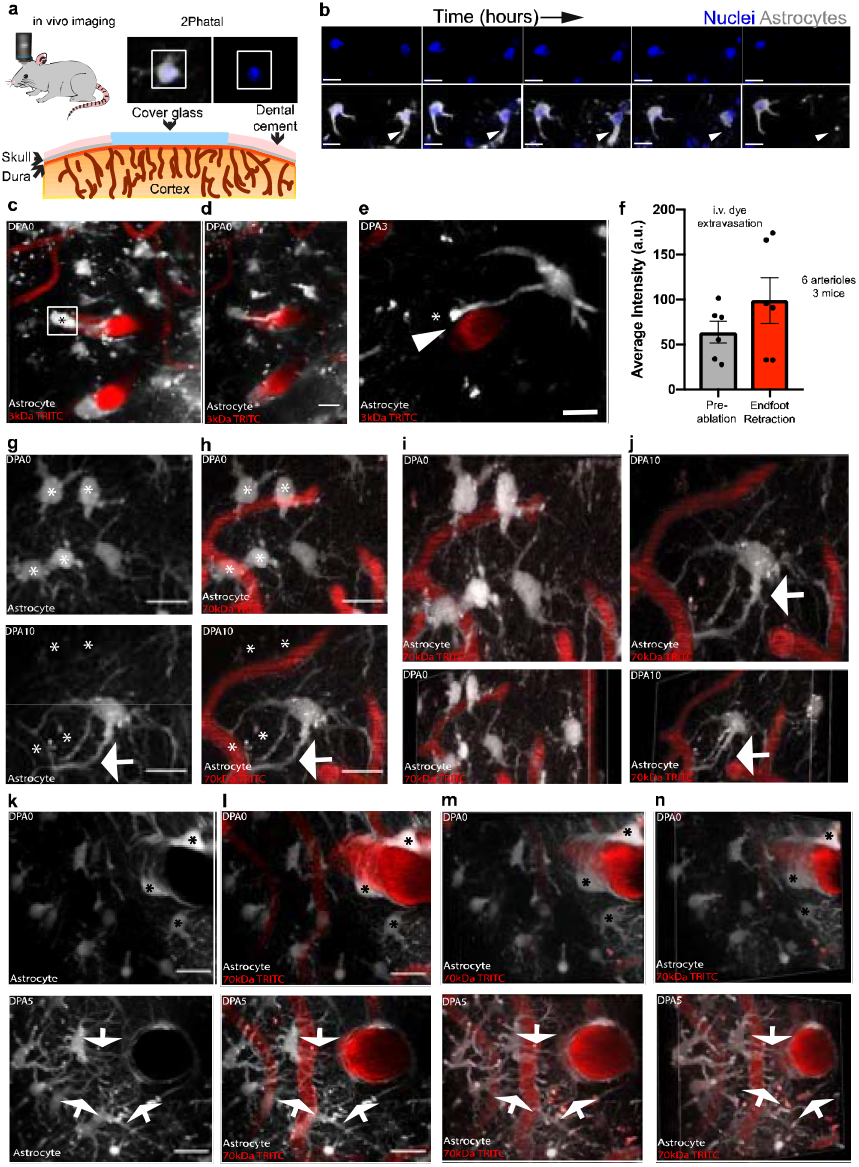
The focal ablation of an astrocyte induces a gliovascular structural plasticity response at all levels of the vascular tree-. To determine if the focal ablation of an astrocyte is sufficient to induce breaches in blood-brain barrier integrity, we utilized the *in vivo* single-cell 2Phatal cellular ablation method. In all images, asterisks indicate the ablated astrocyte(s) and arrows demarcate replacement processes. **a)** Cartoon diagram depicting the experimental approach, including the bath application of Hoechst at the time of surgery and prior to cranial window implantation. By using low laser-power to activate Hoechst, we could **b)** visualize astrocytes in Aldh1l1-eGFP mice up to and beyond removal of their cell body and associated processes, scale bar = 15 μm. **c)** Volumetric reconstruction of an astrocyte at a penetrating arteriole on day post-ablation 0 (dpa0), with **d)** maximum intensity projection of the same field, scale bar = 10 μm, and **e)** maximum intensity projection at dpa5 showing no apparent disruption to blood-brain barrier integrity. Instead, an astrocyte polarizing a process to the vacant vascular location can be seen, scale bar = 10 μm. **f)** Average intensity quantification of 3kDa TRITC extravasation at baseline relative to the moment of endfoot retraction. n = 6 vessels across 3 mice, two-tailed paired end t-test, p < 0.1100. **g)** and **h)** Maximum intensity projection of astrocytes surrounding capillaries at dpa0 (top) and dpa10 (bottom), scale bar = 15 μm. **i)** Volumetric reconstruction at dpa0 of capillary field showing both dorsal (top) and dorsolateral (bottom) views compared to **j)** at dpa10. The arrow indicates replacement processes from neighboring astrocytes. Representative image for n=40 astrocytes/4 mice **k)** and **l)** Maximum intensity projection of an ascending venule at dpa0 (top) and dpa5 (bottom). Arrows represent replacement processes from neighboring astrocytes, scale bar = 20 μm. **m)** Volumetric reconstruction showing dorsal view at dpa0 (top) and dpa5 (bottom). **n)** Same image from the dorsolateral view at dpa0 (top) and dpa5 (bottom). Representative image for n=40 astrocytes/7 mice.

Given that astrocyte endfeet have been reported to cover up to 99% of the entire cerebrovascular surface (34), we then asked if this process occurred at all levels of the vascular tree. To answer this, Alexa Fluor 633 hydrazide was again employed to specifically label arterioles, and Alexa 633 negative vessels larger than 10μm were identified as venules. All vessels smaller than 10μm were identified as capillaries **(Supplementary Figure 1)**. 2Phatal ablation of astrocytes revealed that the replacement of endfeet also occurred at capillaries **(Figure 1g-i)** and venules **(Figure 1k-n)**. **(Figure 1)**. Taken together, these data suggest that focal loss of single astrocytes is sufficient to induce an endfoot replacement response from nearby surrounding astrocytes, regardless of vessel type.

### Replacement endfeet can vasoconstrict precapillary arterioles

Astrocytes have been reported to mediate neurovascular coupling at precapillary arterioles (8). Furthermore, laser-activation of astrocytes has been shown to be a convenient way to probe their contribution to vascular physiology, as it induces a focal rise in intracellular calcium which subsequently leads to the release of vasoactive molecules (35). In order to determine if replacement endfeet have the machinery to perform neurovascular coupling, and thus cause changes in blood vessel diameter, we ablated astrocytes in Aldh1l1-cre x GCaMP5G mice occupying vascular territories on precapillary arterioles. We then laser-activated astrocytes that extended processes to the vacant vascular locations. We consistently observed the laser stimulation triggering an increase in intracellular calcium, immediately followed by a decrease in vessel diameter (**Fig. 2e, i, j and Supplementary video 1**). Compared to original astrocytes, the induced constriction by replacement astrocytes occurred with a similar kinetic profile (**Fig 2m-n**). A higher change in laser-induced original astrocyte intracellular calcium correlated with a higher magnitude of vessel constriction (**Fig. 2k, m and Supplementary video 2**). These data suggest that replacement astrocytes are capable of assuming the original astrocyte’s role in gliovascular coupling, or at a minimum have the ability to release vasoactive molecules, regardless of their existing relationship with a blood vessel.

**Figure 2.**
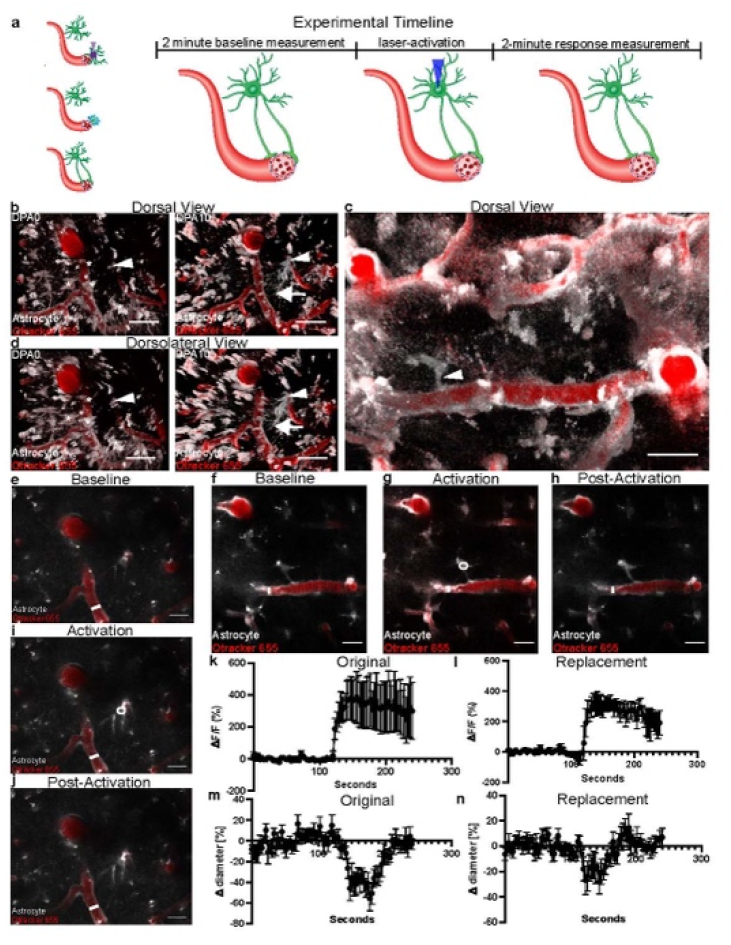
Replacement endfeet vasoconstrict precapillary arterioles-. To determine if replacement endfeet can vasoconstrict precapillary arterioles, we 2Phatal ablated astrocytes in Aldh1l1cre x GCaMP5G mice, targeting astrocytes contacting the first branching capillary segment from an Alexa-633 hydrazide positive penetrating arteriole. **a)** Cartoon depicting experimental paradigm. Upon 2Phatal ablation, astrocytes undergo microglial engulfment, and a new astrocyte subsequently extends processes to reinnervate the vascular vacancy. Replacement astrocytes underwent two minutes of imaging to assess baseline vessel diameter and calcium levels, were then laser-activated, and another two minutes of imaging ensued. **b)** Left-side image is a volumetric reconstruction showing a dorsal view of the field of interest at baseline. The right-side image shows that same field at dpa10. Annotation is as follows: asterisks indicate ablated astrocytes, arrows indicate replacement astrocyte processes, and arrow-heads indicate replacement astrocyte to-be. **c)** and **d)** Volumetric reconstruction depicting a field of interest, with the arrow-head indicating astrocyte chosen for laser-activation, from c) a dorsal view and d) a dorsolateral view of the replacement astrocyte. **e)** Single-optical section of the replacement astrocyte field at baseline, as depicted in b and d. **f)** Single-optical section of the original astrocyte field at baseline, as depicted in c. **g)** Single-optical section showing the original astrocyte and pre-capillary arteriole at the time of laser-activation, followed by **h)** a single-optical section showing return to baseline. **i)** Single-optical section showing the replacement astrocyte and pre-capillary arteriole at time of laser-activation. The precapillary arteriole constricts soon after astrocyte activation. **j)** Single-optical section showing return to baseline. **k)** Quantification of change in fluorescence over baseline fluorescence (Δf/f) for the duration of the experiment in astrocytes originally occupying an appositional vascular location. **l)** Quantification of change in diameter over baseline diameter (Δ diameter) for the duration of the experiment in astrocytes originally occupying an appositional vascular location. **m)** Quantification of change in fluorescence over baseline fluorescence (Δf/f) for the duration of the experiment in replacement astrocytes. **n)** Quantification of change in diameter over baseline diameter (Δ diameter) for the duration of the experiment in replacement astrocytes. n=5 astrocytes over 3 mice for both groups. Scale bars=20 μm

### Endfoot replacement slows with aging

Studies in humans (36), rodents (37–39), and primates (40) indicate that astrocytic morphology in aged astrocytes differs markedly from young astrocytes, and other reports have shown that endfeet actually retract later in life (20). We therefore asked how aging would impact focal endfoot replacement. First, we wanted to confirm that this process remained intact at all levels of the vascular tree, consistent with the observations in young mice. 2Phatal ablation of astrocytes at arterioles, venules, and capillaries revealed that this was indeed the case **(Figure 3a-f).** To determine if aging impacted the fidelity of replacement, we quantified the number of cells extending or growing new processes to the vacant vascular region at each vessel type and found no difference between age groups (**Figure 3g-i**).

**Figure 3.**
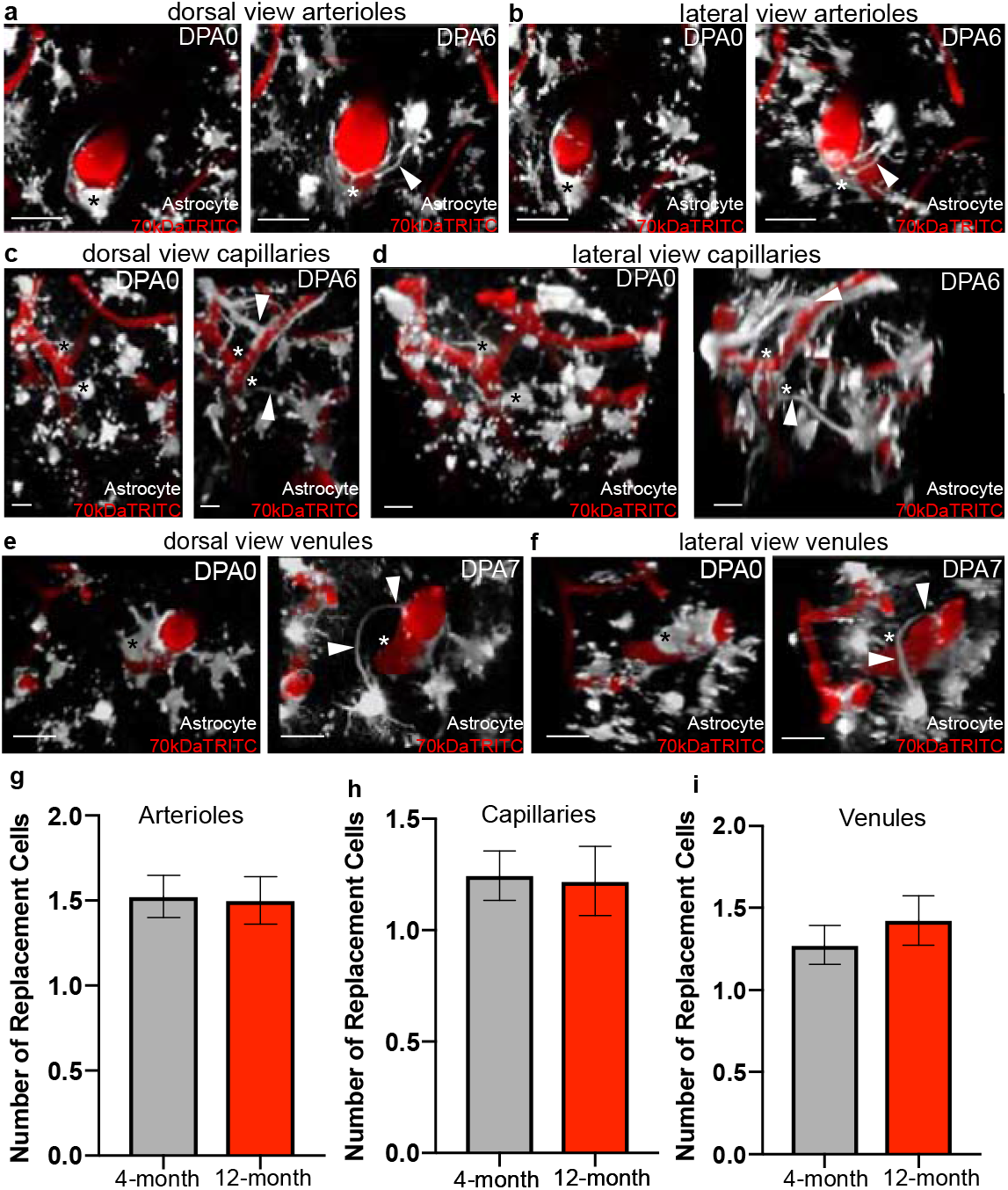
Gliovascular structural plasticity remains intact at levels of the vascular tree in aging-. In order to determine if gliovascular structural plasticity remained intact in aging, we ablated astrocytes in 12-month-old Aldh1l1-eGFP mice making contact with Alexa 633 hydrazide positive penetrating arterioles, capillaries, and venules. We further compared the extent of replacement in old versus young mice as well. In all images, asterisks indicate the ablated astrocyte(s) and arrows demarcate replacement processes. **a)** and **b)** Volumetric reconstruction showing astrocyte associations with a penetrating arteriole at dpa0 (left) and dpa6 (right), from a) a dorsal perspective and b) a dorsolateral perspective. Scale bar=25 μm. **c)** and **d)** Volumetric reconstruction showing the astrocytes and their association with a capillary at dpa0 (left) and dpa6 (right) from c) a dorsal view and d) a dorsolateral view. Scale bar=10 μm. **e)** and **f)** Volumetric reconstruction showing an ascending venule and astrocyte interactions with it at dpa0 (left) and dpa7 (right), from e) a dorsal view and f) a dorsolateral view. Scale bar=20 μm **g)** Average number of replacement astrocytes (the number of cells polarizing processes to vascular vacancies) in 4-versus 12-month old mice at arterioles, Two-tailed Mann-Whitney test, p=0.9465, n=40cells/7 mice for 4-month data, n=40cells/6 mice for 12-month data. **h)** Average number of replacement astrocytes in 4-versus 12-month old mice at capillaries, Two-tailed Mann Whitney test, p=0.6853, n=40 cells/4 mice for both 4- and 12-month old mice. **i)** Average number of replacement astrocytes in 4-versus 12-month old mice at venules, Two-tailed Mann Whitney test, p=0.3060, n=40 cells/7 mice for 4-month old data, n=40cells/5 mice for 12-month old data.

A recent study documented a not only enhanced, but more importantly, prolonged astrogliosis response in aged mice following TBI (41). This ultimately suggest that aging would impact the velocity of an astrocyte response, rather than the extent of it. We therefore next sought to determine if the kinetics of replacement significantly slowed with aging. Engaging in long-term repetitive *in-vivo* imaging revealed that this was indeed the case. 2–4-month-old mice on average had an endfoot replacement event 17 minutes prior to endfoot retraction of the ablated cell **(Figure 4a-b**).In contrast, however, this process was significantly slowed in aged mice to an average replacement time of 112 minutes after endfoot retraction of the ablated cell (**Figure 4c-d, analysis in Figure 4f**). Finally, given that aged animals had vascular vacancies unoccupied for roughly two hours following endfoot retraction of the previously ablated astrocyte, we again aimed to determine if 3kDa TRITC would extravasate at this location. Results revealed this to not be the case **(Supplementary Figure 3)**, which suggests that vascular vacancies unoccupied by astrocytes for this duration of time are not sufficient to disrupt BBB integrity.

**Figure 4.**
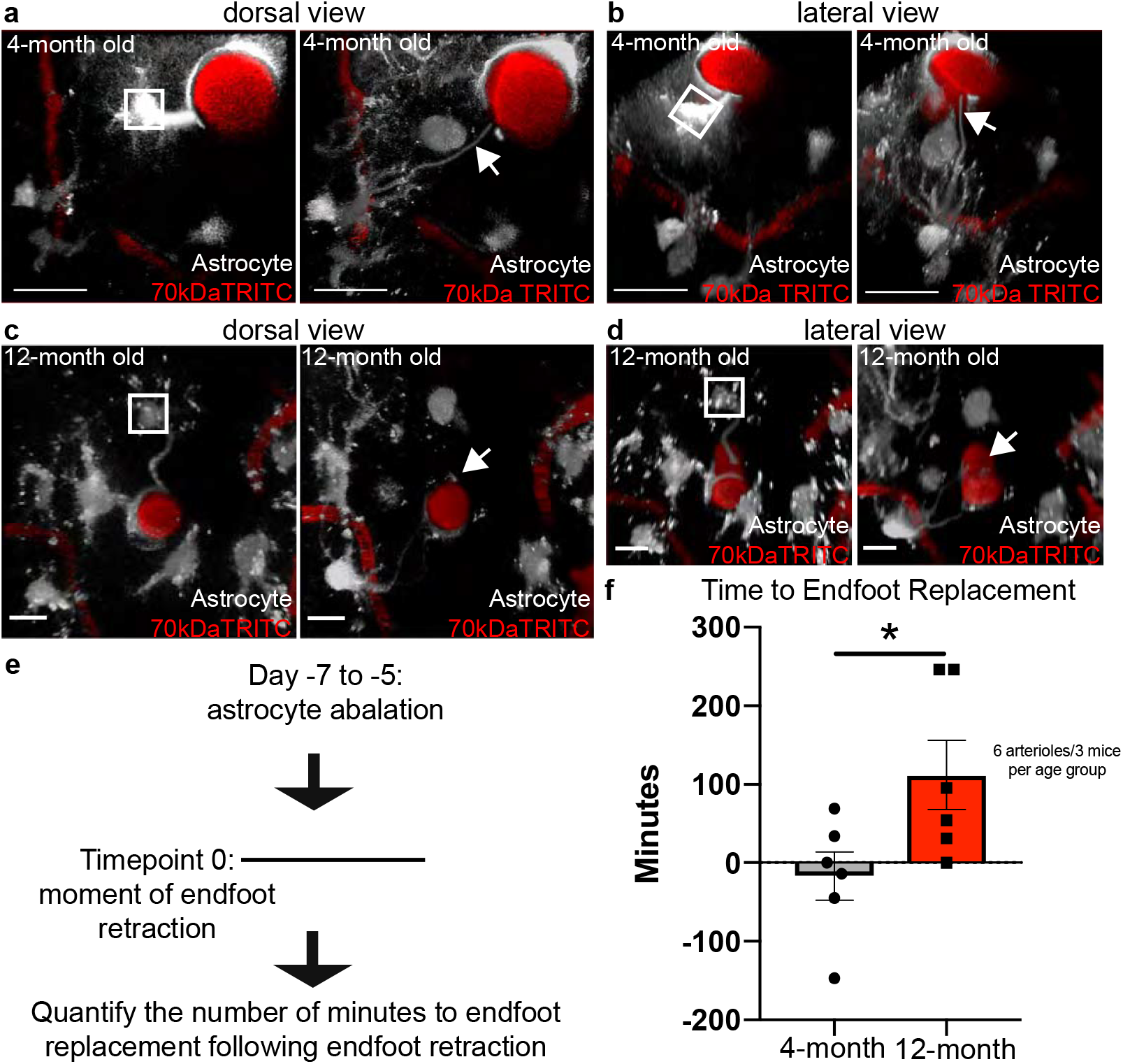
Aging significantly attenuates the kinetics of the gliovascular structural plasticity response-. In order to determine if the kinetics of endfoot replacement significantly slowed as a result of aging, we ablated astrocytes at penetrating arterioles and acquired continual z-stacks to capture the exact time of endfoot replacement. White boxes indicate ablated astrocyte and arrows demarcate replacement processes (as in a and b) or lack thereof (as in c and d). **a)** and **b)** Volumetric reconstruction showing the penetrating arteriole in a 4-month-old mouse at baseline (left) and the exact moment of endfoot retraction (right), from a) a dorsal view and b) dorsolateral view of the same field. Scale bar=25 μm. **c)** and **d)** Volumetric reconstruction showing a penetrating arteriole in a 12-month-old mouse at baseline (left) and the near exact moment of endfoot retraction (right), in c) the dorsal view and d) the dorsolateral view. Scale bar=10 μm e) Schematic illustrating methodology to quantify the number of minutes to endfoot replacement. Right-side images in a-d represent time point zero, or the moment of endfoot retraction. Subsequent images were analyzed to determine the number of minutes until a process occupied an appositional vascular location, as depicted in right-side images of a and b. **f)** Average number of minutes to endfoot replacement in 4-versus 12-month-old mice. n=6 cells/4 mice for both age groups, Two-tailed, unpaired t-test, p<0.0371.

### Pharmacological inhibition of STAT3 phosphorylation via subcutaneous injection of AG490 significantly impairs the endfoot plasticity response

When first questioning which signaling pathways might underlie gliovascular structural plasticity, we noted that the phenotype observed in Figure 1 at venules and capillaries appeared markedly like reactive astrogliosis. This suggested that molecules previously shown to be mediators of gliosis would be valid candidates to explore. The phosphorylation of the signal transducer and activator of transcription 3 (pSTAT3) molecule by janus kinase 2 (JAK2) has long been known to underlie astrogliosis, as its pharmacological and/or genetic inhibition results in a significantly dampened gliotic response. This is evidenced by a significant reduction in glial fibrillary acidic protein (GFAP), which is considered to be a marker of reactive astrogliosis (42–43). To test the hypothesis that pSTAT3 is necessary for focal endfoot replacement, we subcutaneously injected the JAK2 inhibitor AG490 over a seven-day period **(Figure 5A)**. Comparing the volume of eGFP signal in astrocyte processes post-ablation in AG490 to vehicle-injected control animals revealed a significantly attenuated response (**Figure 5b-f),** suggesting that pSTAT3 is indeed a necessary arbiter of the focal endfoot replacement response. We further wanted to assess if any perturbations in BBB integrity ensued at a penetrating arteriole after an attenuated plasticity response, and as our previous results would suggest, this was not the case (**Figure 5g-h**).

**Figure 5.**
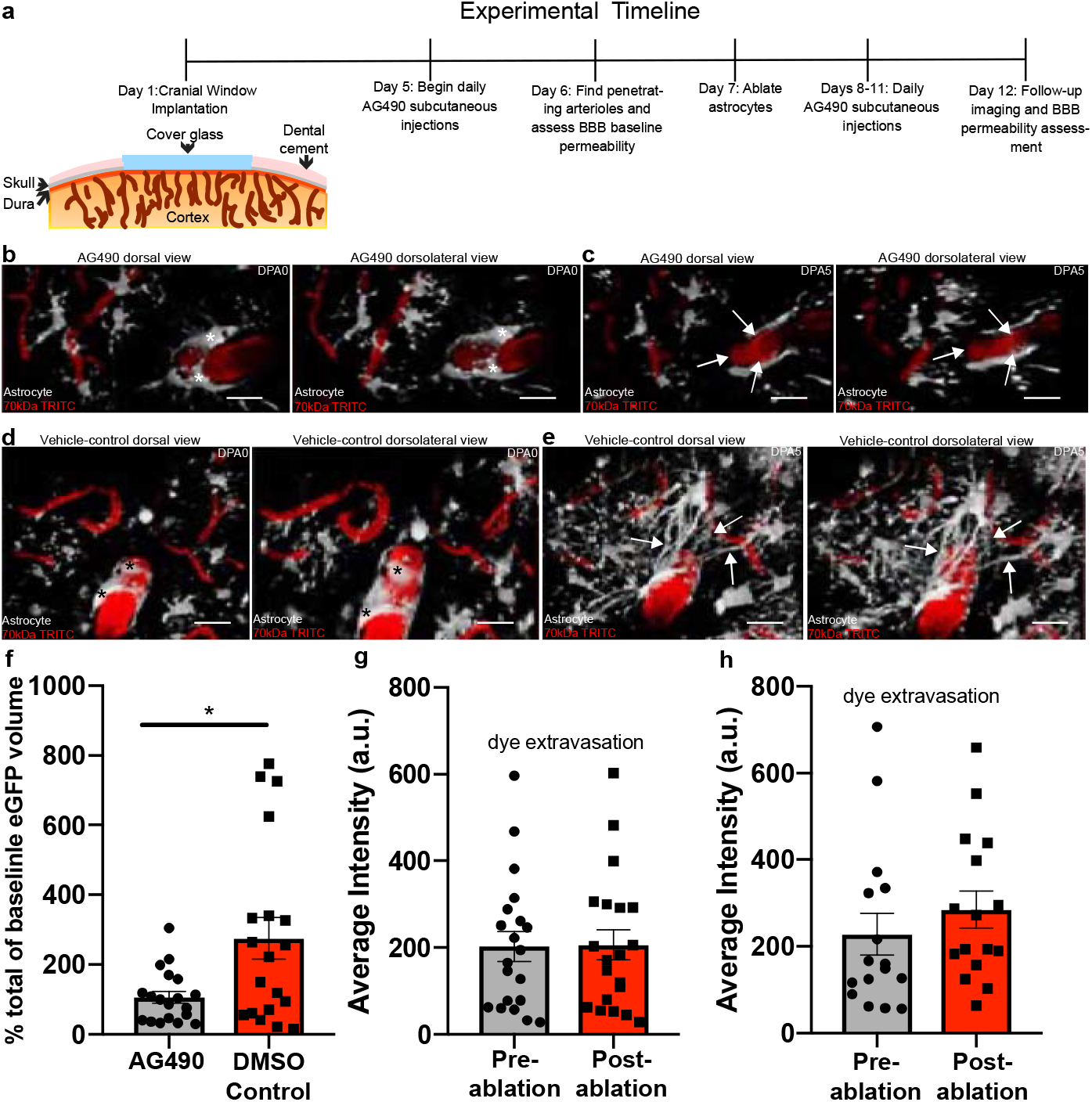
The pharmacological inhibition of EGFR/pSTAT3 significantly reduces the volumes of replacement astrocytes post-ablation-. In order to determine if EGFR and pSTAT3 is necessary for gliovascular structural plasticity, we subcutaneously injected AG490. In all images, asterisks indicate ablated astrocyte(s) and arrows demarcate replacement processes, or lack thereof. **a)** Schematic illustrating the experimental timeline. **b)** and **c)** Volumetric reconstructions showing the dorsal view (left) and dorsolateral view (right) of astrocytes and their interactions with a penetrating arteriole in AG490-injected mice at b) dpa0 and c) dpa5. Scale bar=20 μm **d)** and **e)** Volumetric reconstructions showing the dorsal view (left) and dorsolateral view (right) of astrocytes interacting with a penetrating arteriole in DMSO vehicle-injected control mice at d) dpa0 and e) dpa5. Scale bar=20 μm. **f)** Bar graph comparing percent increase in eGFP volume at dpa5 relative to baseline in AG490-injected mice relative to DMSO-injected control, n=19 penetrating arterioles/4 mice, Two-tailed Mann-Whitney test, p=0.0497. **g)** Bar graph comparing the average intensity of 3kDa TRITC extravasation pre-ablation versus post-ablation, n=20 penetrating arterioles/4 mice, Two-tailed Wilcoxin matched pairs signed-rank test, p=0.6215. **h)** DMSO vehicle-injected control group, n=16 penetrating arteriole/4 mice, Two-tailed Wilcoxin matched pairs signed-rank test, p=0.1167.

Given that we only attenuated an overall increase in astrocyte volume following one dosage per day of AG490, we wanted to determine how BBB integrity might be impacted if we significantly reduced overall astrocyte volume at arterioles post-ablation when increasing AG490 dosage to three times per day. Results revealed that, though we were able to significantly reduce the gliovascular structural plasticity response, we were not able to completely abolish it. Furthermore, even in locations along penetrating arterioles that were severely stripped of endfoot coverage, the BBB remained intact (**Supplementary Figure 4b-e**). Taken together, these results suggest that a significant loss of endfoot coverage is not sufficient to disrupt BBB integrity.

### 2Phatal as a model of focal endfoot replacement following astrocyte loss post-transient photothrombotic stroke

We initially turned to 2Phatal to model a loss of endfoot coverage on the underlying vasculature based off what had we had previously characterized in two disease conditions (23–24). 2Phatal, however, results in complete loss of an astrocyte cell body and associated processes rather than just an endfoot; we therefore wanted to determine if 2Phatal might more closely model other disease conditions. Given that 2Phatal presumably triggers apoptosis through ROS-induced DNA damage (30), we searched for reports on any disease conditions marked by astrocyte cell death at the vascular interface due to ROS-induced DNA damage. Ischemiareperfusion is a condition whereby blood-flow is restored to a vessel following an ischemic insult due to vessel occlusion, and astrocytes have been reported to be just as sensitive as neurons to reperfusion-induced apoptosis following restoration of blood flow (44).

We therefore set out to model this condition *in vivo* by utilizing the Rose Bengal photothrombosis stroke model. Rose Bengal is a light-sensitive dye that, upon encountering a green laser, undergoes nucleation and forms a clot. Successful clot formation can be visualized by a dark area forming in the vessel, indicative of red blood cell accumulation, and intense fluorescence due to dye accumulation above that dark mass (**Figure 6d-f**) (45). Critically, this method allowed us to focally occlude penetrating arterioles and thereby examine if a structural plasticity response would occur following focal loss of astrocyte-vascular coverage. In all instances of successful vessel occlusion, reperfusion occurred within 1 hour (data not shown), making this more akin to a transient ischemic attack rather than a stroke. We subsequently looked in regions below the plane of dye nucleation for signs of astrocyte cell death at the vascular interface. Cell death was indeed observed, and in the days following, surrounding astrocytes reached out to that vacant vascular location **(Figure 6g-I, compare to baseline images in Figure 6a-c)**. This data suggests that the 2Phatal ablation of astrocytes at vascular interfaces could potentially be thought of as a model for focal loss of endfoot coverage due to reperfusion-induced apoptosis. Given that stroke does lead to neural injury and subsequent astrogliosis, these data further support the notion that gliovascular plasticity is a focal gliosis response aimed at ensuring continual vascular coverage by astrocytes.

**Figure 6.**
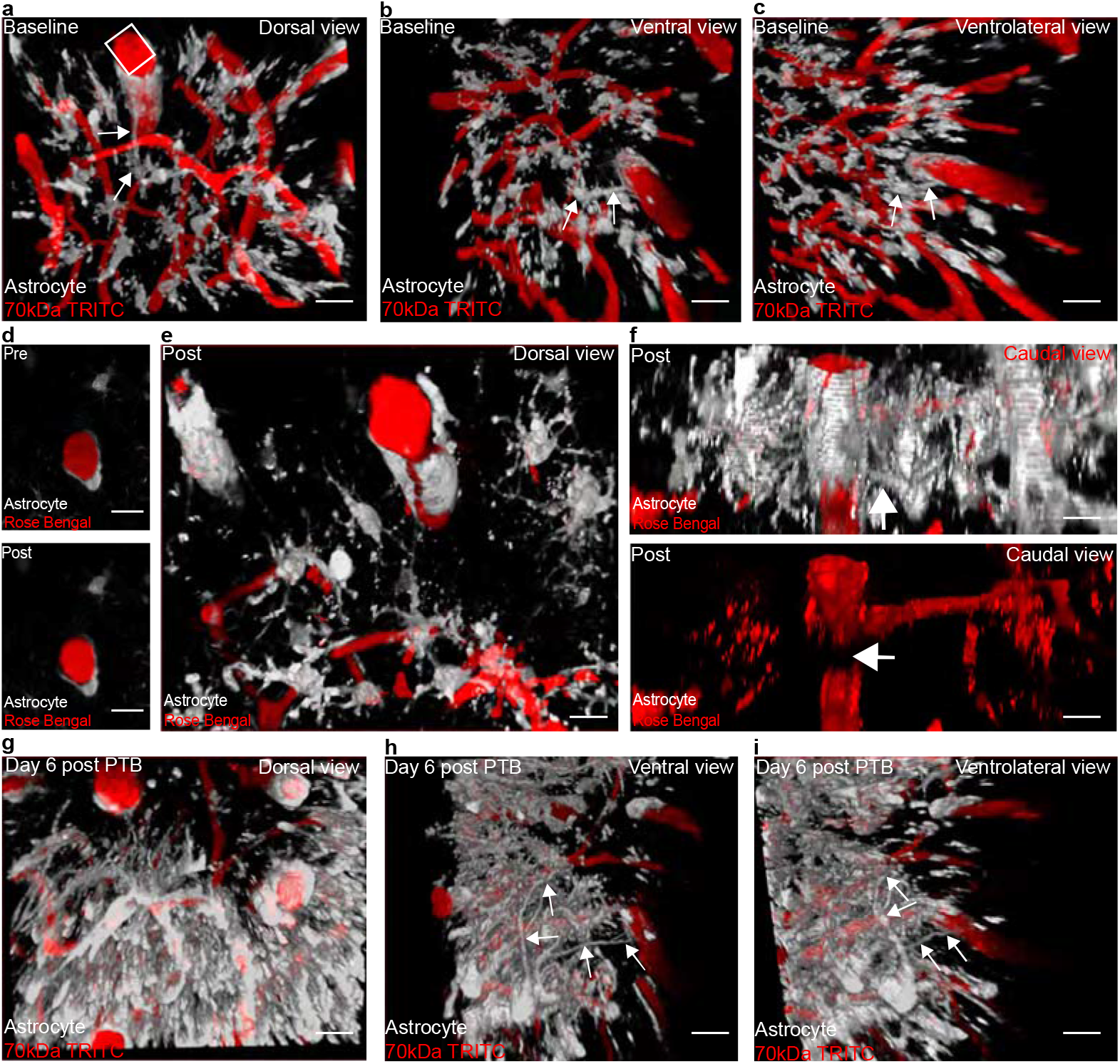
Gliovascular structural plasticity occurs following reperfusion post-focal photothrombotic stroke-. In order to determine if gliovascular structural plasticity occurs following loss of astrocyte-vascular coverage due to CNS insult, we turned to the Rose Bengal focal photothrombotic stroke method. Arrows in all images demarcate astrocyte processes either at baseline (as in a, b, c, and f) or replacement processes six days following reperfusion from focal-stroke (as in h-i). **a)** through **c)** Volumetric reconstruction showing a penetrating arteriole at baseline from a) dorsal view, b) a ventral view, and c) a ventrolateral view. **d)** Single optical section showing a penetrating arteriole following Rose Bengal injection (top) and immediately after laser-induced dye nucleation (bottom). **e)** and **f)** Volumetric reconstruction showing a penetrating arteriole immediately after Rose Bengal laser-induced dye nucleation from e) a dorsal view and f) a lateral view of that same field showing the penetrating arteriole after Rose Bengal injection (top) and (bottom) immediately after dye nucleation. The arrow in the bottom image demarcates where RBC buildup has occurred. **g)** Volumetric reconstruction showing a dorsal view of the previously occluded vessel six days later. Note that reperfusion occurred within the hour following Rose Bengal dye nucleation (data not shown) **h)** Volumetric reconstruction of the same field but from a ventral view and **i)** ventrolateral view. n=5 arterioles/5 mice. Scale bar in a-i=20 μm

## Discussion

Previous studies suggest that a number of nervous system insults and diseases present with impaired gliovascular interactions and even BBB disruption. Here we set out to determine if focal ablation of single astrocytes is sufficient to compromise BBB integrity focally. We had previously shown that focal endfoot separation due to invading glioma cells resulted in extravasation of various molecular weight dextran dyes and significant losses in tight junction proteins zonula-occludens-1 and Claudin 5 (23). By employing the 2Phatal ablation technique to induce single-cell apoptosis, we found that focal loss of an astrocyte did not compromise bloodbrain barrier integrity **(Figure 1e-f)**, but instead reliably induced a plastic response whereby surrounding astrocytes reach out their processes to fill the vascular vacancy left by the ablated astrocyte **(Figure 1e)**. Moreover, the BBB remained intact in conditions where the vasculature was vacant for prolonged periods of time **(Supplementary Figure 3c)** or was almost entirely stripped of endfoot coverage **(Supplementary Figure 4).**

These results are interesting given a recent study (31) demonstrating the necessity of astrocytes in maintaining BBB integrity. These prior findings were obtained using a sparser and more permanent astrocyte ablation and relied on extravasation of ~1kDa Cadaverine as a marker for BBB disruption. Unfortunately, in our studies we found this dye flawed in its ability to discriminate between normal and abnormal BBB function, since we observed baseline leakage in control mice. **(Supplementary Figure 2a)**. Another recent study (46) that used albumin, a reliable extravasation marker, showed that an astrocyte-specific connexin-30 and −43 double knockout resulted in both swollen astrocytic endfeet and impaired BBB integrity. However, this was most prevalent in deep brain structures such as the striatum and basal ganglia rather than cortical regions, which is very similar to what was demonstrated upon deletion of astrocytespecific laminins (47). Note that care must be taken to account for changes in pericyte support, given that pericyte-deficient mouse models have been shown to alter astrocyte properties (48); therefore, loss of astrocytes may have affected pericyte coverage and thus indirectly altered BBB integrity. To avoid a confounding contribution of pericyte dysfunction to BBB integrity, we exclusively studied penetrating arterioles where pericytes are absent.

While it is conceivable that the role of astrocytes in maintaining BBB integrity may be region-specific, which aligns with evidence supporting astrocyte functional heterogeneity in the brain (49), we believe that our data points to the rapid plasticity or repair response by neighboring astrocytes as the primary reason that vessel function and BBB integrity are unaffected by the loss of a single or few astrocytes. This is in excellent agreement with a recent study that focally ablated pericytes, which similarly did not damage BBB integrity-but did result in a comparable plasticity response (25). Yet, global pericyte-deficient mouse models have been clearly shown to perturb the BBB (48). Unlike focal pericyte ablation, which resulted in an absence of pericyte-capillary coverage for days, focal ablation of astrocytes in our hands only results in a lapse of endfoot coverage for minutes to hours. Indeed, we were surprised to find that the reinnervation of the blood vessel by replacement astrocytes typically preceded complete retraction of the lesioned cell by a few minutes. Given that the half-life for the tight junction protein ZO-1 is 5.2 hours in MDCK cells (50), and 90 minutes for claudin-5 (51)’ it is likely that we were unable to strip a vessel of endfoot contact long enough to breach the barrier, assuming focal ablation is sufficient to do so. It is further possible that the pharmacological inhibition of STAT3 prevented BBB breakdown following astrocyte ablation, as other studies have documented restoration of BBB integrity following prevention of STAT3 activation via inhibition of JAK (52). Future studies focally ablating astrocytes should aim to do so in conditions where the replacement of endfeet is either stalled for longer periods or completely abolished. Alternatively, such studies may find-as was the case in focal pericyte ablation studies-that a more permanent focal ablation of astrocytes is not sufficient to perturb BBB integrity.

Beyond the BBB, we also built off a previous study that documented endfoot plasticity (53) and further determined that it occurs at all levels of the vascular tree **(Figure 1g-n)**. Furthermore, this process begins even prior to the ablated astrocyte having its corpse entirely removed in young mice, whereas aging significantly slows down the swift kinetics of replacement **(Figure 4c-f)**. These results are interesting given our observations of endfoot plasticity following transient ischemic attack (TIA) **(Figure 6).** It is known that the probability of stroke occurrence following TIA increases with aging (54), and presumably any reduction of or lapse in endfoot coverage could affect vessel recovery following ischemic onset-potentially contributing to this increased stroke probability.

Finally, inhibiting the phosphorylation of STAT3 by JAK2 via AG490 injection revealed pSTAT3 to be an essential regulator of the focal endfoot plasticity response. Though many studies have documented AG490 inhibition of JAK2-mediated phosphorylation of STAT3 (42, 55–56), AG490 also inhibits the epidermal growth factor receptor (EGFR). Given that pharmacological inhibition of EGFR with the specific inhibitor PD168393 also attenuated reactive astrogliosis following spinal cord injury (57), these results point to focal endfoot replacement response as being dependent on genes classically associated with astrogliosis (58).

Taken together, this is the first study to characterize gliovascular structural plasticity at all levels of the vascular tree, its physiological relevance to blood flow and the BBB, and its alterations in aging. These findings add to the burgeoning literature regarding the complexities of basic astrocyte biology and the role of the astrocytes in supporting the cerebrovasculature.

## Acknowledgements

This work was supported by NIH grants 1R01AG065836, 1R01CA227149, and 1RO1-NS082851

---

**Supplementary Figure 1.**
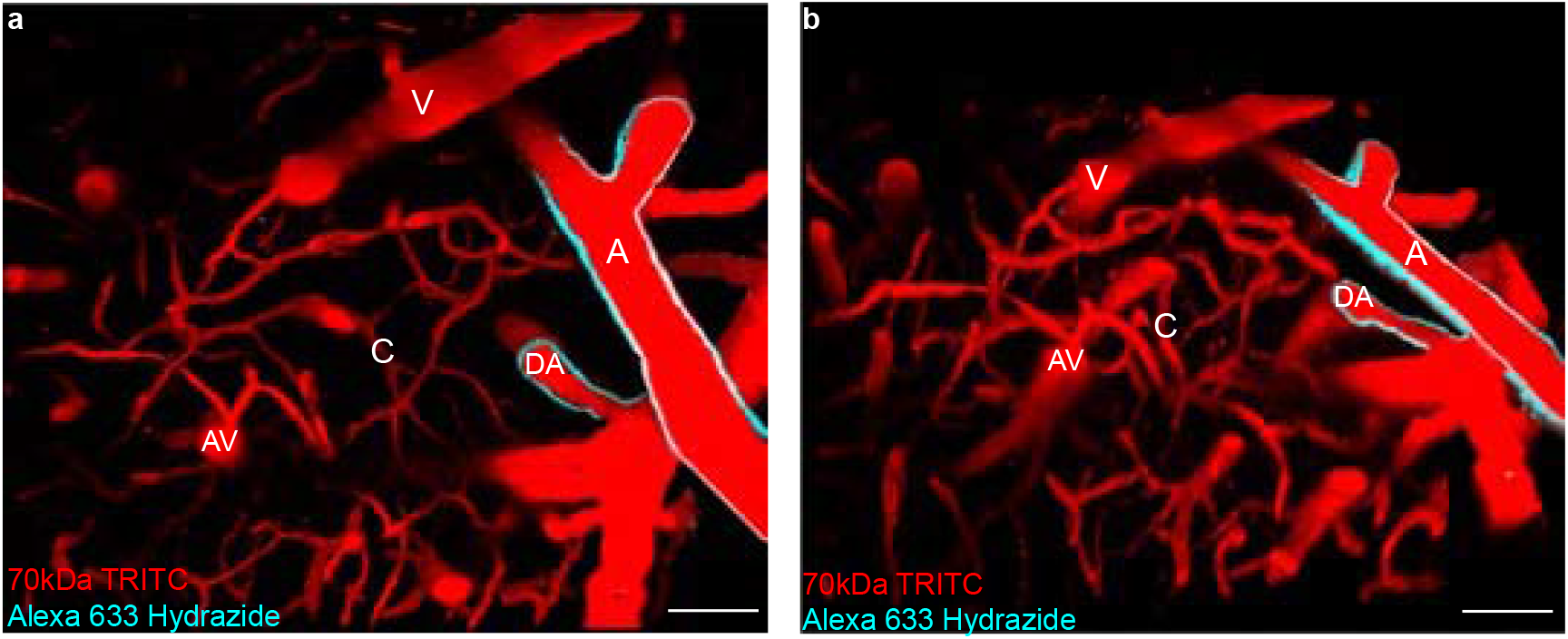
Alexa 633 hydrazide positively selects for arterioles-. In order to determine which vessel type we were ablating astrocytes at, we employed the use of Alexa 633 hydrazide to: positively select for arterioles and penetrating arterioles; size exclusion; and negative selection of venules, ascending venules, and capillaries (<10 μm). Volumetric reconstruction showing **a)** a dorsal view of all zones of the vascular tree and **b)** a dorsolateral view of the same field. Scale bar=25 μm. A-arteriole, DA-descending arteriole, V-venule, AV-ascending venule, C-capillary.

**Supplementary Figure 2.**
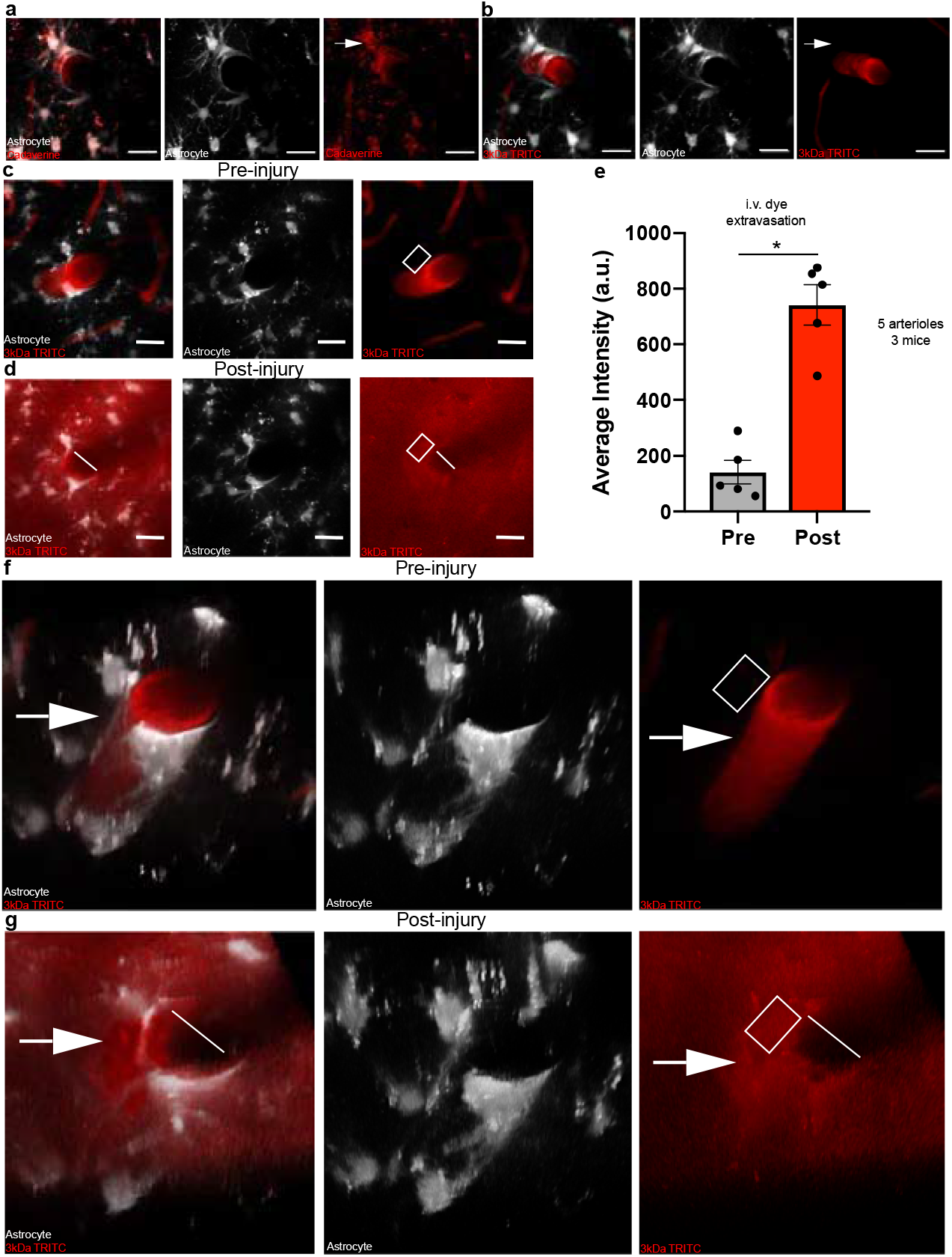
Extravasation of 3kDa TRITC can be reliably detected *in vivo*-. To measure breaches in BBB integrity, we employed the use of small molecular weight dyes and determined through laser-induced vessel injury that they can be reliably detected *in vivo*. **a)** Maximum intensity projections showing cadaverine extravasation at baseline in Aldh1l1-eGFP mice (left, middle, and right). **b)** Maximum intensity projections showing that 3kDa TRITC does not extravasate at baseline in Aldh1l1-eGFP mice (left, middle, and right). Scale bar in a-b=20 μm **c)** Maximum intensity projections showing a penetrating arteriole at baseline without apparent 3kDa TRITC extravasation relative as opposed to **d)** obvious 3kDa TRITC dye extravasation following vessel injury. The white line indicates where the laser scanned across the vessel lumen, and the white box indicates where measurements were taken. Scale bar in c and d=20 μm. **e)** Comparison of the average intensity of 3kDa TRITC extravasation pre-vessel injury relative to post-injury. n=5 arterioles/3mice, Two-tailed paired t-test, p<0.0021. **f-g)** Volumetric reconstructions showing a dorsolateral view of the same images displayed in c and d.

**Supplementary Figure 3.**
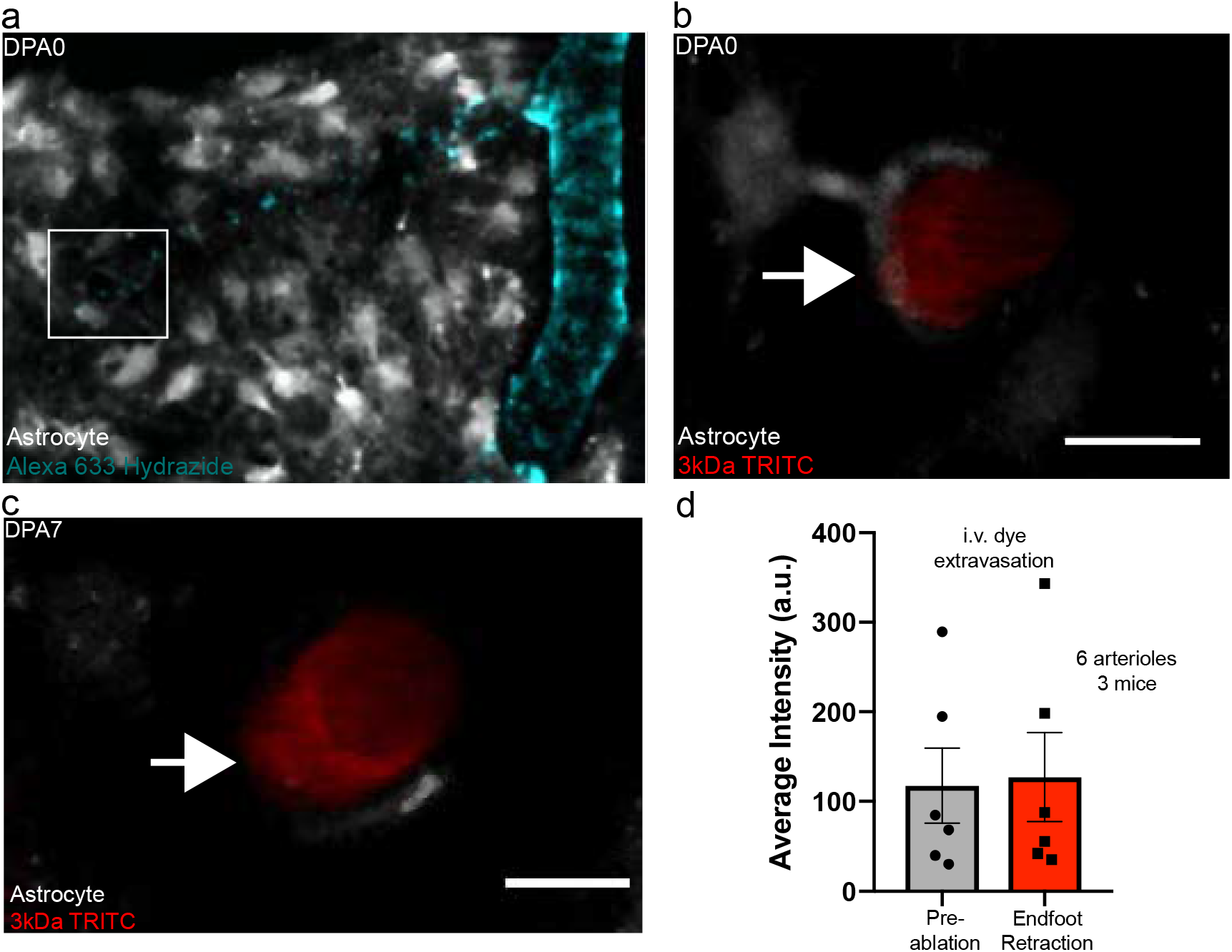
Blood-brain barrier integrity is not compromised in 12-month-old mice. **a)** Maximum intensity projection showing an Alexa-633 hydrazide positive arteriole in a 12-month-old Aldh1l1-EGFP mouse. White square indicates a penetrating portion of a branch from that arteriole, shown **b)** at a higher magnification at baseline, and **c)** at the same magnification at the exact moment of endfoot retraction on dpa7. Scale bar in b and c = 10 μm. **d)** Comparison of the average intensity of 3kDa TRITC at baseline, prior to astrocyte ablation, relative to post-ablation. n=6 arterioles/3mice, Two-tailed paired t-test, p<0.3785.

**Supplementary Figure 4.**
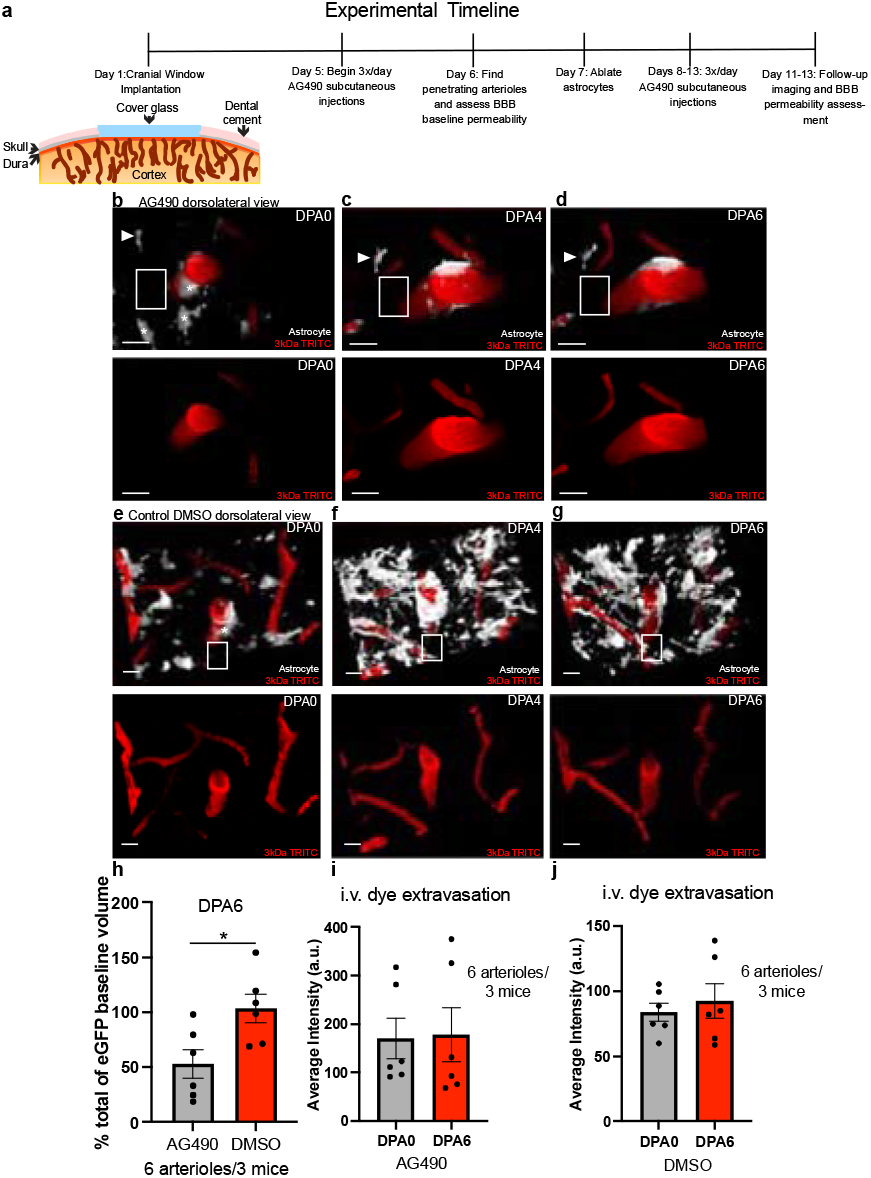
Significant reductions in astrocyte volume at penetrating arterioles post-ablation is sufficient to maintain BBB integrity-. In order to determine how BBB integrity would be impacted following a significant reduction in endfoot coverage post-astrocyte ablation, we adopted the experimental timeline outlined in **a)**. Following cranial window implantation, mice received 3 doses of AG490 daily beginning at day five postoperation. This injection continued through post-operation day 13. After ablating astrocytes on day seven post-operation, follow-up imaging would occur to assess endfoot plasticity volume for three days (days four to six post-ablation). **b)** Volumetric reconstruction at baseline depicting a dorsolateral view of the astrocyte-vascular landscape (top) or just vascular landscape (bottom) in AG490-injected mice. **c)** Same field as b, but day 4 post-ablation, and **d)** at day 6 post-ablation. Arrow-head indicates a reference astrocyte. Asterisks indicate ablated astrocytes. Scale bar in b-d= 20 μm. **e)** Volumetric reconstruction at baseline depicting a dorsolateral view of the astrocyte-vascular landscape (top) or just vascular landscape (bottom) in DMSO control-injected mice. Asterisk indicates the ablated astrocyte. **f)** Same field as e, but at day 4 post-ablation, and **g)** at day 6 post-ablation. **h)** Quantification of the percent change in astrocyte volume at dpa6 relative to dpa0 along arteriole locations where astrocytes were ablated. n=6 arterioles over 3 mice. Two-tailed unpaired t-test, p< 0.0213. **i)** Average intensity of 3kDa TRITC dye extravasation along arteriole locations where astrocytes were ablated at dpa6, relative to baseline in AG490-injected mice. n=6 arterioles/3 mice. Two-tailed paired t-test. P<0.4104. **j)** Average intensity of 3kDa TRITC dye extravasation along arteriole locations where astrocytes were ablated at dpa6, relative to baseline in DMSO-injected control mice. n=6 arterioles/3 mice. Two-tailed paired t-test. p< 0.5861. White boxes in b-e represent locations of ROI placement for BBB measurements. Scale bar in e-g=10 μm.

## Notes

### Competing Interest Statement

The authors have declared no competing interest.

